# *Dystrophin* interacts with *Msp300* to Regulate Myonuclear Positioning and Microtubule Organization

**DOI:** 10.1101/2024.11.07.622444

**Authors:** Jorel R. Padilla, Yunshu Qiu, Grace Aleck, Lillie Ferreria, Sharon Wu, William Gibbons, Torrey Mandigo, Eric S. Folker

## Abstract

During *Drosophila* myognesis, myonuclei are actively moved during embryogenesis, and their spacing is maintained through an anchoring mechanism in the fully differentiated myofiber. While we have identified microtubule associated proteins, motors, and nuclear envelope proteins that regulate myonuclear spacing, the developmental time during which each gene functions has not been tested. Here we have identified a *Dystrophin* as required only for the maintenance of myonuclear spacing. Furthermore, we demonstrate that *Dystrophin* genetically interacts with the KASH-domain protein *Msp300* to maintain myonuclear spacing. Mechanistically, both *Dystrophin* and *Msp300* regulate microtubule organization. Specifically, in animals with disrupted expression of both *Dystrophin* and *Msp300*, microtubule colocalization with sarcomeres is reduced. Taken altogether, these data indicate that the peripheral membrane protein Dystrophin, and the outer nuclear membrane protein Msp300, together regulate the organization of the microtubule network which then acts as an anchor to restrict myonuclear movement in contractile myofibers. These data are consistent with growing evidence that myonuclear movement and myonuclear spacing are critical to muscle development, muscle function, and muscle repair and provide a mechanism to connect disparate muscle diseases.

**Summary Statement:** Here we show that *Dystrophin* is required to maintain the spacing of nuclei in differentiated myofibers. Furthermore, *Dystrophin* achieves this function via a genetic interaction with *Msp300* which regulates microtubule organization.

## Introduction

The regulated positioning of the nucleus is conserved across all eukaryotes (Calero-Cuenca et al., 2018; Gundersen and Worman, 2013; Osorio and Gomes, 2014), but the mechanisms that drive nuclear positioning are often cell-specific. The multinucleated myofiber provides a unique opportunity to identify mechanisms that regulate the spatial organization of nuclei because the myonuclei undergo multiple movements during development that are themselves independently regulated. Although there are some variations between species, the basic principles of myonuclear spacing are conserved from *Drosophila* to mammals. First, the nucleus of a recently fused myoblast is moved to the center of the developing myofiber, the myotube. Then, the myonuclei are dispersed throughout the myotube and spaced evenly apart from one another. Finally, after the myotube has differentiated into the contractile myofiber, myonuclei are anchored in place to maintain their spacing during contraction (Bone and Starr, 2016; Bruusgaard et al., 2006; Roman and Gomes, 2018; Wilson and Holzbaur, 2015). The proper spacing of myonuclei has been linked to muscle function, while the active movement of myonuclei has been linked to repair (Metzger et al., 2012; Roman et al., 2021). Critically, that active myonuclear movement in the myotube and maintenance of myonuclear position in the myofiber occur at different times, in cells with significantly different structures, suggests that the molecular mechanisms regulating movement and maintenance may be different.

The links between development, muscle function, and myonuclear spacing are best understood in *Drosophila melanogaster* which allows for efficient visualization of the developing myotube in embryos, and the differentiated myofiber in larva (Azevedo et al., 2016). The cellular structure of the myofiber is conserved between *Drosophila* and humans but the breadth of genetic tools and the absence of complex bundling of cells makes *Drosophila* significantly more amenable to cell biological analysis. Furthermore, during normal repair in mammalian myofibers, nuclei move and occupy different positions making it difficult to discern mispositioned nuclei from newly incorporated nuclei *en route* to their proper position (Chargé and Rudnicki, 2004). The absence of muscle repair in *Drosophila* ensures any misplaced nuclei are not merely an indication of ongoing repair.

The benefits of *Drosophila* have been leveraged to determine that myonuclear positioning is dependent on microtubules, microtubule motors, traditional microtubule associated proteins (MAPs), and various nuclear envelope (NE) associated proteins (Collins et al., 2021; Dialynas et al., 2010; Elhanany-Tamir et al., 2012; Folker et al., 2014; Mandigo et al., 2019; Metzger et al., 2012; Rosen et al., 2019). Specifically, each previously described mechanism requires the Linker of Nucleoskeleton and Cytoskeleton (LINC) complex which directly links the cytoskeleton and the nuclear lamina (Elhanany-Tamir et al., 2012; Mandigo et al., 2019). Critically, the genes that regulate this connection between the myonucleus and the cytoskeleton have been implicated in Emery–Dreifuss muscular dystrophy (EDMD) (Attali et al., 2009; Bione et al., 1994; Bonne et al., 1999; Muntoni et al., 2006).

More broadly, mispositioned myonuclei is associated with many muscle diseases, including Duchenne muscular dystrophy DMD (Folker & Baylies, 2013; Wang et al., 2000). However, it is unknown whether mispositioned myonuclei are the result of a disrupted spacing mechanism, ongoing muscle repair, or both. DMD is caused by a mutation in the gene *Dystrophin* which encodes a protein that is localized to the Dystroglycan complex (DGC) located at the sarcolemma (Rentschler et al., 1999). *Dystrophin* stabilizes the sarcolemma through the DGC, and disruption of this process contributes to the deterioration of muscle in DMD patients (Watkins et al., 1988). However, the microtubule network of myofibers is disorganized in the mouse model for DMD, the *mdx* mouse (Khairallah et al., 2012; Oddoux et al., 2019). Furthermore, Dystrophin has a characterized microtubule binding function is crucial for muscle function (Belanto et al., 2014).

Here, we have characterized the effect of *Dystrophin* on myonuclear spacing. Using temporal and tissue specific RNAi, we identify *Dystrophin* as required specifically for the maintenance of myonuclear spacing. This is the first example a gene that only impacts myonuclear anchoring, with no apparent role for the active movement of myonuclei in the myotube. Furthermore, we describe a genetic interaction between *Dystrophin* and the LINC complex component *Msp300* in regulating myonuclear spacing. Finally, we found that *Dystrophin* and *Msp300* genetically interact to regulate the colocalization of microtubules and F-actin thin filaments in differentiated myofibers. These data demonstrate a novel mechanism for *Dystrophin*-dependent muscle development. Furthermore, these data link DMD and EDMD related genes supporting the hypothesis that even myonuclear spacing is a fundamental feature of properly functioning muscle.

## Results

### The maintenance of myonuclear spacing is *Dystrophin*-dependent

Previous studies have shown that myonuclear spacing and microtubule organization are disrupted in *mdx* mice which express a truncated and nonfunctional *Dystrophin* gene (Liu et al., 2020; Oddoux et al., 2019). However, these studies only examined differentiated myofibers and did not determine whether *Dystrophin* contributed to the active translocation of myonuclei during muscle development, the maintenance of myonuclear spacing, or both. To determine the developmental timing of *Dystrophin* function with respect to myonuclear spacing we expressed RNAi against *Dystrophin* during distinct developmental stages using the GAL4-UAS system, and measured the spacing of myonuclei in differentiated larval myofibers as previously described (Collins & Mandigo et al., 2017; Mandigo et al., 2019). Twist-GAL4-dependent expression of RNAi against *Dystrophin* during active myonuclear movement did not affect myonuclear spacing (Fig. 1 A,B). However, Mhc-GAL4-dependent expression of RNAi against *Dystrophin* after myonuclei had achieved their final spacing caused significant clustering of myonuclei (Fig. 1 C,D). Twist-GAL4-dependent expression of *Dystrophin* RNAi results in a transient depletion of the protein with *Dystrophin* expression being restored during late embryonic development. To determine whether embryonic nuclear position was *Dystrophin*-dependent, we measured the position of myonuclei in developing embryonic myotubes as previously described (Collins et al., 2021; Folker et al., 2012) and found that Twist-GAL4-dependent expression of *Dystrophin* RNAi did not affect myonuclear position during embryonic development (Fig. 1 E,F,G). To support the RNAi data, we examined the effects of a null mutation in *Dystrophin*. The homozygous mutation was lethal but the muscles from animals heterozygous for the null mutation had defects in myonuclear spacing (Fig. 2).

**Figure 1:**
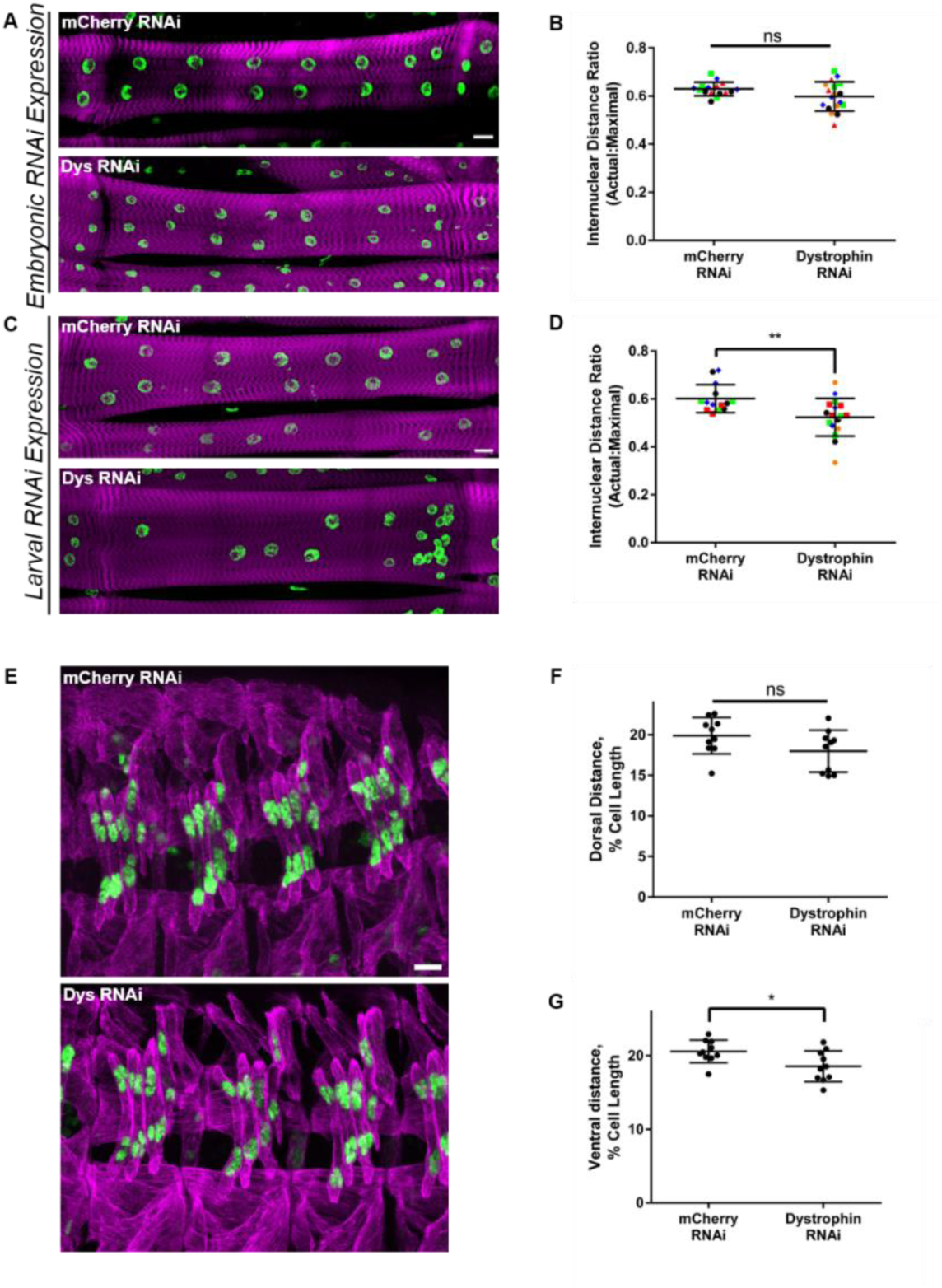
*Dystrophin* is necessary to maintain, but not establish myonuclear spacing. **(A)** Fluorescence images of VL3 muscles expressing RNAi against indicated gene under control of the embryonic mesodermal-specific driver Twist-GAL4. Magenta, phalloidin/muscles; green, Hoechst/nuclei. Scale bar = 25μm. (**B)** Internuclear distance ratio in larval muscles expressing the indicated RNAi. Each point is one myofiber. Myofibers from the same animal are the same shape and color. Data compared by t-test. Four myofibers were analyzed from four animals across 2 biological replicates. **(C)** Fluorescence images of VL3 muscles expressing RNAi against the indicated gene under control of the larval muscle-specific driver Mhc-GAL4. Magenta, phalloidin/muscles; green, Hoechst/nuclei. **(D)** Internuclear distance ratio in larval muscles expressing the indicated RNAi. Each point is one myofiber. Myofibers from the same animal are the same shape and color. Data compared by t-test. Four myofibers were analyzed from four animals across 2 biological replicates. P < 0.05 **(E)** Immunofluorescence images of LT muscles in stage 16 embryos expressing RNAi against indicated gene under control of the embryonic mesodermal-specific driver Twist-GAL4. Magenta, tropomyosin/muscles; green, apRed/nuclei. Scale bar = 10μm. **(F)** Average distance from the dorsal end of the muscle to the nearest nucleus. **(G)** Average distance from the ventral end of the muscle to the nearest nucleus. F and G: Each point is the average of LT1, LT2, LT3 from three hemisegments in a single embryo. Data compared by t-test. 10 Animals were analyzed across two separate biological replicates. P < 0.01

**Figure 2:**
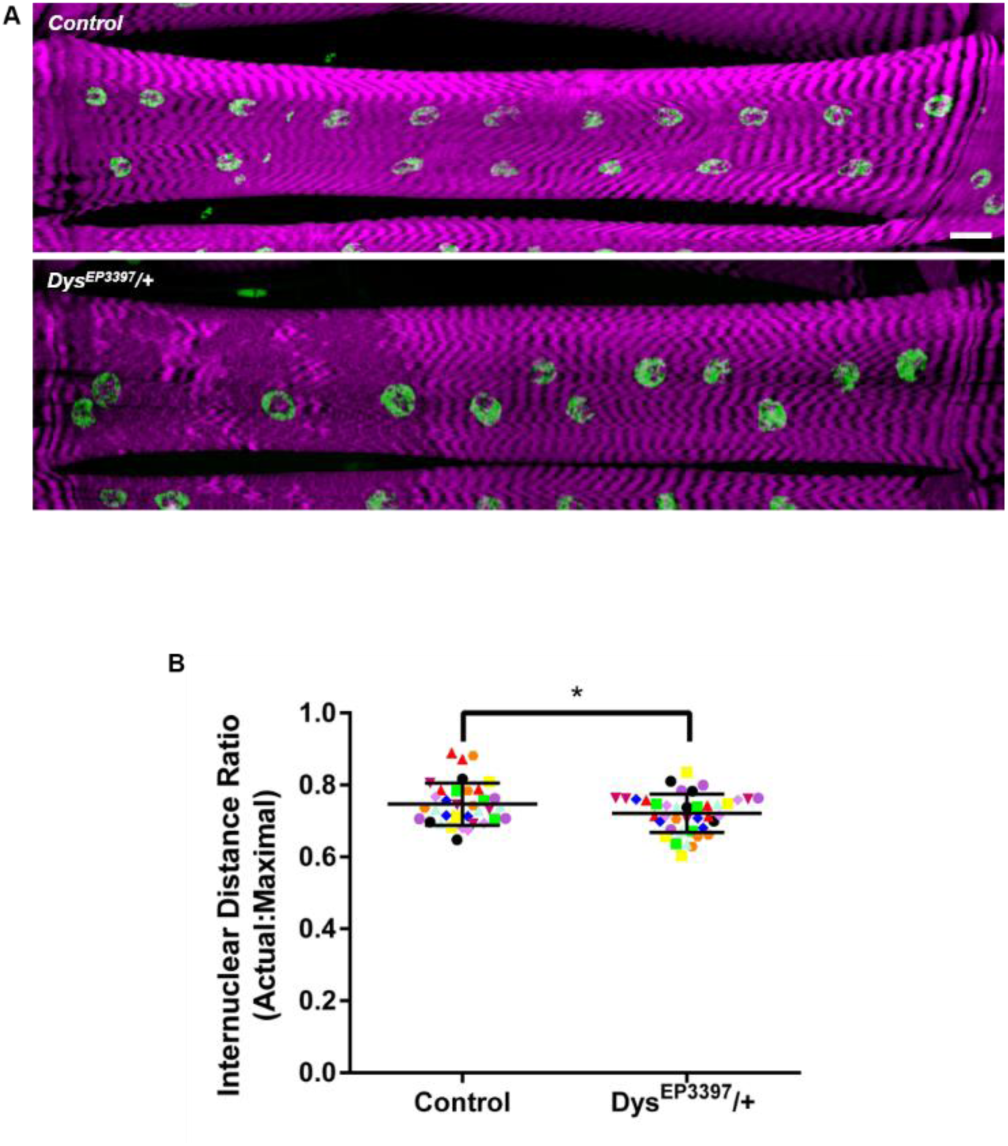
A *Dystrophin* null mutation disrupts myonuclear spacing. **(A)** Fluorescence images of VL3 muscles with indicated genotypes. Magenta, phalloidin/muscles; green, Hoechst/nuclei. Scale bar = 25μm. **(B)** Internuclear distance ratio in larval muscles from animals with the indicated genotype. Each point is one myofiber. Myofibers from the same animal are the same shape and color. Data compared by t-test. Four myofibers were analyzed from four animals across two biological replicates. P < 0.05

Recent work showed that in animals homozygous for RNAi insertions at the *attp40* docking site, *Msp300* function was impaired and resulted in myonuclear positioning defects (van der Graaf et al., 2022). Myonuclear spacing was normal in animals heterozygous for insertions at the *attp40* docking site. Consistent with these data, animals heterozygous for both *Dystrophin* RNAi and Twist-GAL4 had normal myonuclear positioning indicating that heterozygocity for the *Dystrophin* RNAi insertion at the *attp40* site does not cause a myonuclear positioning defect. To test whether the insertion of Mhc-GAL4 was sufficient to affect myonuclear spacing, we measured the spacing of myonuclei in larvae homozygous for the Mhc-GAL4 insertion and found that myonuclear spacing was similar to controls (Fig. S1). These data indicate that Mhc-GAL4 is not inserted in a position to impair *Msp300* function and suggests that the myonuclear positioning defects are due to disrupted expression of *Dystrophin*. From here forward, all RNAi experiments will utilize Mhc-GAL4 dependent expression so the GAL4 will not be specified. Altogether, these experiments demonstrate that *Dystrophin* is not necessary for active myonuclear movement, but is necessary to maintain myonuclear spacing in differentiated myofibers.

*Dystrophin* functions through the multi-protein Dystroglycan Complex (DGC) to regulate sarcolemmal integrity, a function that when compromised contributes to the onset and progression of Duchenne muscular dystrophy (Watkins et al., 1988). Therefore, we tested whether the DGC is also necessary to maintain myonuclear spacing. We expressed RNAi against several DGC components, and found that myonuclear spacing was not dependent on any DGC complex proteins (Fig. 3 A,B). Therefore, *Dystrophin* regulates myonuclear spacing independently of the DGC complex with which *Dystrophin* has well-described interactions.

**Figure 3:**
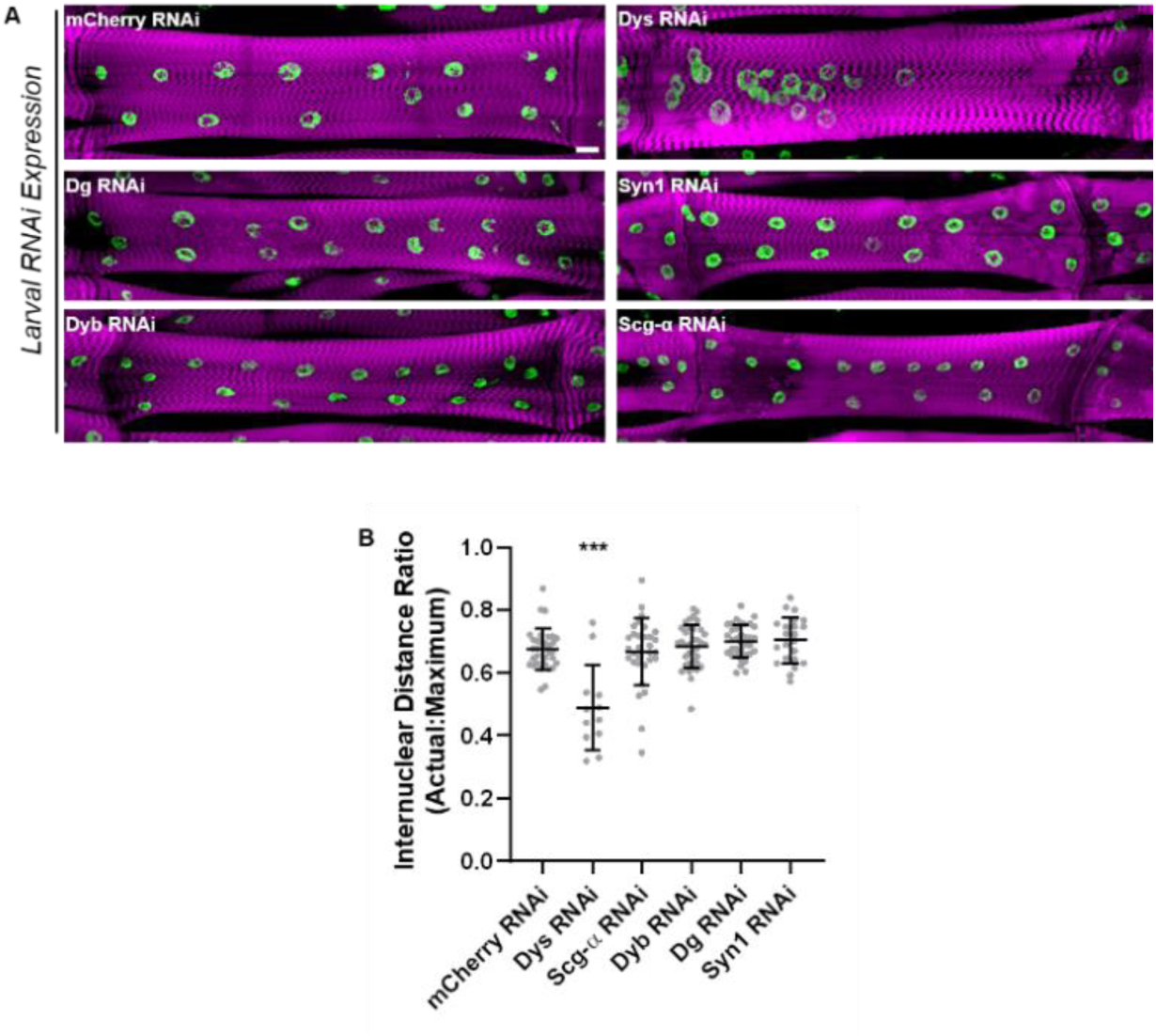
Individual DGC proteins do not regulate myonuclear spacing. **(A)** Fluorescence images of VL3 muscles expressing RNAi against the indicated gene under control of the larval muscle-specific driver Mhc-GAL4. Magenta, phalloidin/muscles; green, Hoechst/nuclei. Scale bar = 25μm. **(B)** Internuclear distance ratio in larval muscles expressing the indicated RNAi. Each point is one myofiber. Data compared by ANOVA with multiple comparisons. Four myofibers were analyzed from four animals across two biological replicates. P < 0.001

### *Dystrophin* regulates the organization of microtubules in myofibers

We and others have shown that myonuclear spacing is regulated by microtubule associated proteins and microtubule motors in *Drosophila* and in mammalian culture systems (Cadot et al., 2012; Collins et al., 2021; Folker et al., 2014; Metzger et al., 2012; Wilson & Holzbaur, 2015). Although microtubule organization is disrupted in the myofibers from *mdx* mice (Khairallah et al., 2012; Oddoux et al., 2013; Oddoux et al., 2019), whether *Dystrophin* regulates microtubule organization in *Drosophila* and whether the effects on microtubule organization contribute to maintaining nuclear spacing have not been tested. In *Drosophila* myofibers, microtubules form a ring around the individual myonuclei and emanate out from each myonucleus (Collins et al., 2021; Metzger et al., 2012; Oddoux et al., 2013). We found that in animals that were either heterozygous for a null mutation in *Dystrophin* (Fig. 4A) or expressed *Dystrophin* RNAi, there were several abnormal features of microtubule organization. First, the intensity of the microtubule ring around each myonucleus was reduced compared to controls (Fig. 4 A,C). Second, the microtubules often looped back towards the myonuclei, rather than extending away from nuclei as seen in controls (Fig. 4A; arrowheads). In addition to these shared phenotypes, the mutation and the RNAi had different effects on microtubule organization near and distant from the myonucleus. *Dystrophin* RNAi reduced the number of nuclei that had a clearly defined nucleus associated ring of microtubules and caused a flattening of the decay in tubulin immunofluorescence intensity distant from the nucleus. The heterozygous mutant was similar to controls in both measurements (Fig. 4C). The increased disruption of microtubule organization is consistent with the more dramatic myonuclear spacing phenotype in animals expressing *Dystrophin* RNAi (Fig. 1 C,D) compared to the heterozygous mutants (Fig. 2 A,B).

**Figure 4:**
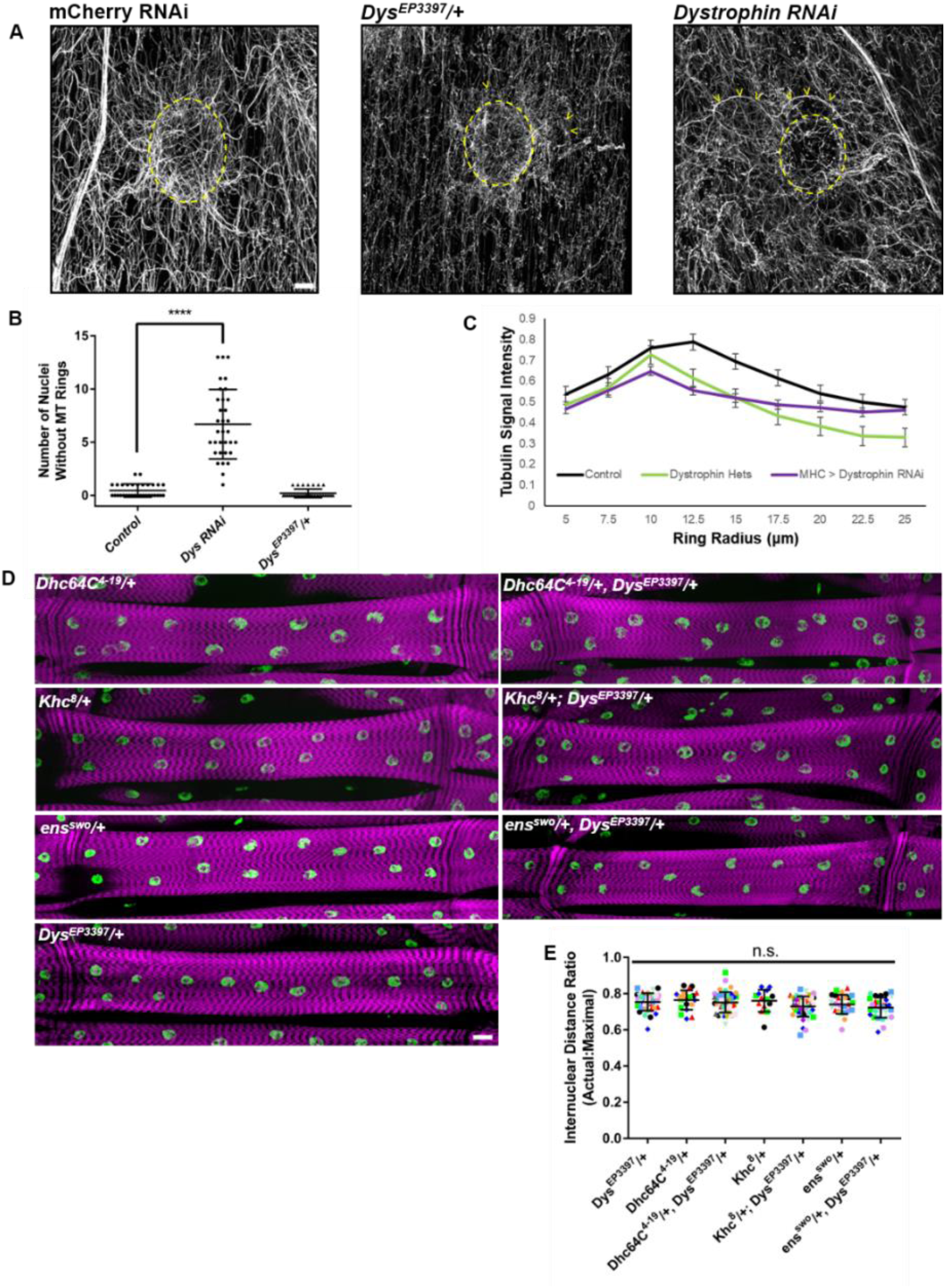
*Dystrophin* regulates microtubule organization in *Drosophila* myofibers. **(A)** Immunofluorescence AiryScan images of the microtubules surrounding a nucleus in VL3 muscles with indicated genotypes. The edge of the myonucleus is marked with a dashed yellow line in each image. Yellow arrowheads point towards microtubules looping back towards the myonucleus. RNAi driven by larval-specific Mhc-GAL4. Gray, α-tubulin. Scale bar = 5μm. **(B)** Graph showing the number of myonuclei without rings in control animals and in animals expressing an RNAi against *Dystrophin.* RNAi driven by larval-specific Mhc-GAL4. Data compared by t-test. Four myofibers were analyzed from four animals across two biological replicates. P < 0.0001 **(C)** Graph showing changes in α-tubulin immunofluorescence intensity as a function of distance from the myonucleus in indicated genotypes. 0 indicates the center of the myonucleus. The peak intensity corresponds to the cytoplasmic side of the myonuclear envelope. Six nuclei from two animals across two biological replicates were analyzed. **(D)** Fluorescence images of VL3 muscles from animals with indicated genotypes. Magenta, phalloidin/muscles; green, Hoechst/nuclei. Scale bar = 25μm. **(E)** Internuclear distance ratio in larval muscles of indicated genotypes. Each point is one myofiber. Myofibers from the same animal are the same shape and color. Data compared by ANOVA with multiple comparisons. Four myofibers were analyzed from four animals across two biological replicates.

Because *Dystrophin* regulates both the maintenance of myonuclear spacing and microtubule organization (Figs. 1 C,D, 3 A,B,C), and nuclear spacing is microtubule-dependent (Bruusgaard et al., 2006; Folker et al., 2014; Metzger et al., 2012; Oddoux et al., 2013, 2019; Wilson & Holzbaur, 2012), we tested whether *Dystrophin* genetically interacts with microtubule associated proteins (MAPs) and microtubule motors to regulate myonuclear spacing by examining double heterozygotes. Critically, we have previously used double-heterozygous screens to place genes in distinct pathways that separately regulate myonuclear positioning (Folker et al., 2012, 2014). We examined animals that were doubly heterozygous for null mutations in *Dystrophin* (*Dys^EP3397^*) and each of cytoplasmic *Dynein* (*Dhc64C^4-19^*), *Kinesin* (*Khc^8^*), and *Ensconsin* (*ens^swo^*, *Drosophila MAP7*), all of which regulate myonuclear spacing (Folker et al., 2012; Metzger et al., 2012). Surprisingly, *Dystrophin* did not genetically interact with any of these factors to regulate myonuclear spacing (Fig. 4 D,E). Therefore, while *Dystrophin* regulates the maintenance of myonuclear spacing and microtubule organization, it does not function with previously identified MAPs and motors that regulate myonuclear spacing.

### *Dystrophin* interacts with the nesprin homolog *Msp300* to regulate the maintenance of myonuclear spacing

In addition to being microtubule-dependent, myonuclear spacing requires several genes that encode nuclear envelope proteins (Collins & Mandigo et al., 2017; Collins et al., 2021; Elhanany-Tamir et al., 2012; Gimpel et al., 2017; Mandigo et al., 2019; Metzger et al., 2012; Wilson & Holzbaur, 2015). Therefore, we tested if there was a genetic interaction between *Dystrophin* and NE-associated genes known to regulate myonuclear spacing. We examined myonuclear spacing from animals that were doubly heterozygous for a null allele of *Dystrophin* (*Dys^EP3397^*) and a null allele each of *Msp300 and Klar* which encode KASH (Klarsicht, Anc, and Syne Homology) domain proteins*, Koi* which encodes a SUN (Sad1p, Unc-84) domain protein*, and Ote* and *Bocks* which encode Emerin homologs (Fig. 5A). Myonuclear spacing in all animals that were doubly heterozygous for *Dys^EP3397^* and either *Klar^1^*, *Koi^EY03560^*, *Ote^B279^*, *Ote^DB^*, or *Bocks^DP01391^* was similar to individual heterozygotes (Fig. 5B). However, animals that were doubly heterozygous for *Dys^EP3397^* and *Msp300^compl^* had a decrease in the evenness of myonuclear spacing when compared to individual heterozygotes (Fig. 5 A,B). These data indicate that *Dystrophin* genetically interacts specifically with *Msp300* to regulate the maintenance of myonuclear spacing.

**Figure 5:**
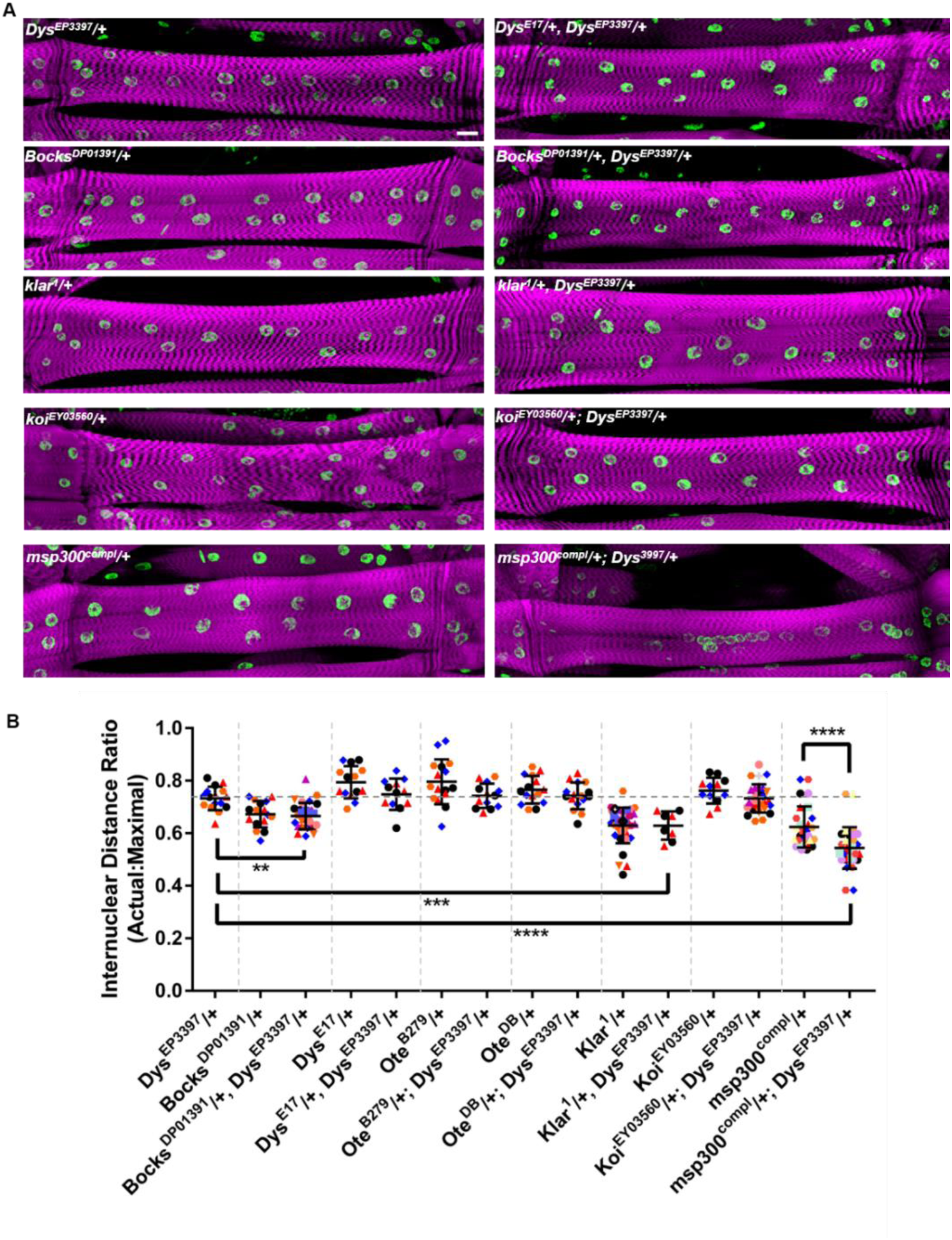
*Dystrophin* and *Msp300* genetically interact to regulate myonuclear spacing. **(A)** Fluorescence images of VL3 muscles with indicated genotypes. Magenta, phalloidin/muscles; green, Hoechst/nuclei. Scale bar = 25μm. **(B)** Internuclear distance ratio in larval muscles from animals with the indicated genotype. Each point is one myofiber. Myofibers from the same animal are the same shape and color. Dashed vertical lines separate experimental comparison groups. The horizontal line indicates the mean internuclear distance ratio in controls. Data compared by ANOVA with multiple comparisons. At least three myofibers were analyzed from three animals across two biological replicates. P < 0.0001

### *Dystrophin* and *Msp300* are necessary for the organization of microtubules in larval myofibers

Having demonstrated that *Dystrophin* and *Msp300* genetically interact to regulate myonuclear spacing, we aimed to determine whether this interaction was functioning through the organization of the microtubule cytoskeleton. Using AiryScan imaging we imaged larval myofibers stained for Dystrophin, F-actin, and tubulin, and found that the Dystrophin signal and the F-actin signal are mostly anticorrelated (Fig. 6 A,C). Interestingly, we also observed that microtubules were more likely to colocalize with F-actin than with Dystrophin (Fig. 6 C,D).

**Figure 6:**
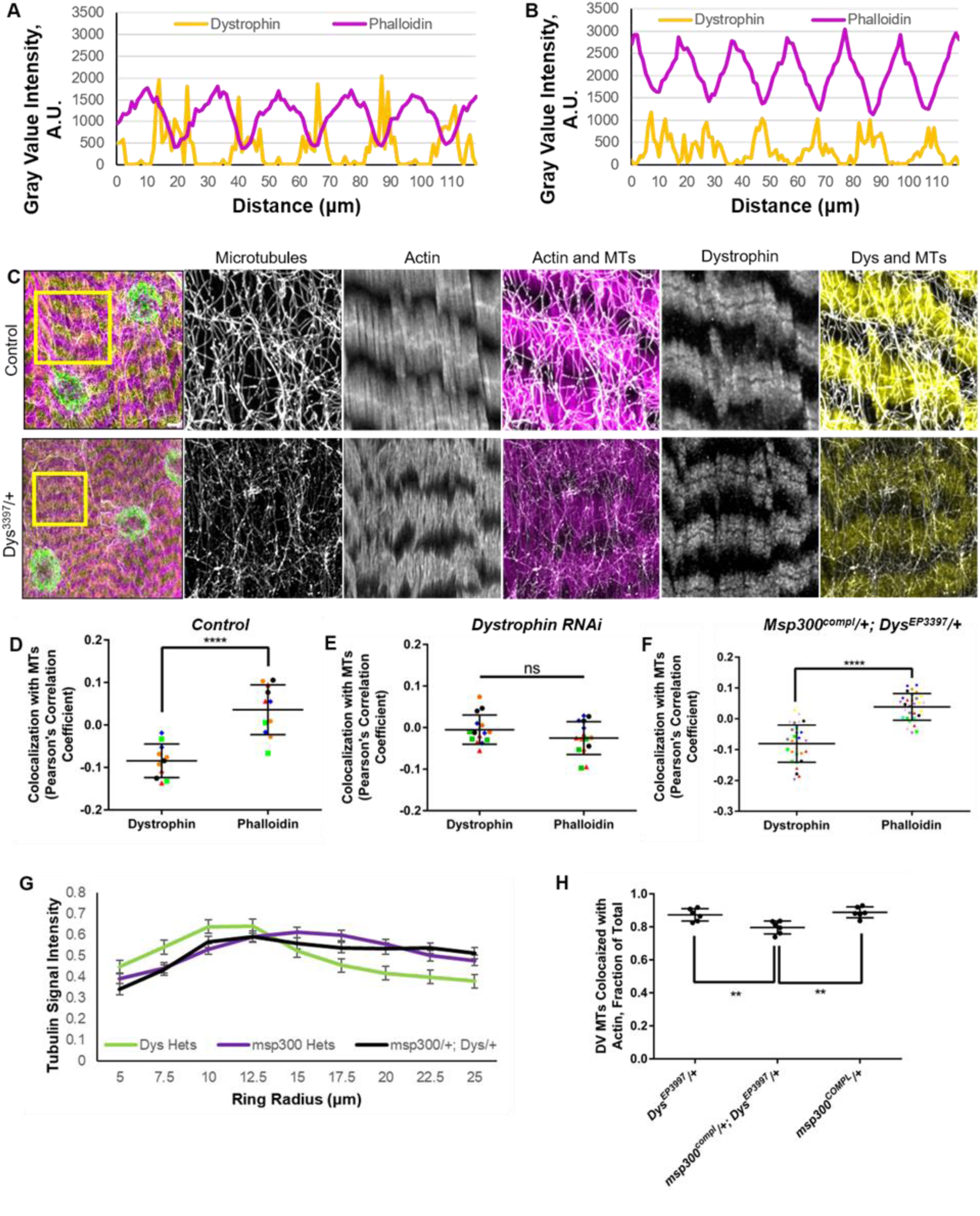
*Dystrophin* is necessary for microtubule colocalization with actin. **(A)** Graph showing the intensity of Dystrophin and phalloidin staining as a function of position on the anterior-posterior axis of a single control myofiber. **(B)** Graph showing the intensity of Dystrophin and phalloidin staining as a function of position on the anterior-posterior axis of a single myofiber expressing RNAi against *Dystrophin* under control of the larval-muscle specific driver Mhc-GAL4. **(C)** Immunofluorescence AiryScan image of a region of a VL3 muscle from a control animal and an animal heterozygous for a null allele of *Dystrophin.* Magenta, phalloidin/muscles; yellow, Dystrophin; gray, α-tubulin; green, Hoechst/nuclei. Yellow boxes indicate regions shown at higher magnification in images to the right. Scale bar = 5μm. **(D)** Pearson’s correlation coefficient describing microtubule colocalization with either Dystrophin or Phalloidin (actin) in a control animal. Each point is the correlation from one X by Y region in a single muscle. Regions from the same animal are the same shape and color. Data compared by t-test. Two regions were analyzed in five animals across two biological replicates. P < 0.0001 **(E)** Pearson’s correlation coefficient describing microtubule colocalization with either Dystrophin or Phalloidin (actin) in an animal expressing *Dystrophin* RNAi under control of the larval-specific Mhc-GAL4 driver. Each point is one X by Y region in a single muscle. Regions from the same animal are the same shape and color. Data compared by t-test. Three regions were analyzed in five animals across two biological replicates. **(F)** Pearson’s correlation coefficient describing microtubule colocalization with either Dystrophin or Phalloidin (actin) in an animal doubly heterozygous for null alleles of *Dystrophin* and *Msp300*. Each point is one X by Y region in a single muscle. Regions from the same animal are the same shape and color. Data compared by t-test. Three regions were analyzed in seven animals across two biological replicates. P < 0.0001 **(G)** Graph showing changes in α-tubulin immunofluorescence intensity as a function of distance from the myonucleus in indicated genotypes. 0 indicates the center of the myonucleus. The peak intensity corresponds to the cytoplasmic side of the myonuclear envelope. Six nuclei from two animals across two biological replicates were analyzed. **(H)** Graph showing the changes in percent of dorsal-ventral microtubules which align with actin between animals doubly heterozygous for null alleles of *Msp300* and *Dystrophin* and animals singly heterozygous for both. Data compared by ANOVA with multiple comparisons. Six images from three animals across two biological replicates. P < 0.01

Because *Dystrophin* has known actin and microtubule binding domains, we tested whether *Dystrophin* was necessary for the microtubule-actin colocalization (Belanto et al., 2014; Fealey et al., 2018). In animals that expressed RNAi against *Dystrophin*, microtubules did not show a preferential colocalization with F-actin (Fig. 6 C,D,E). To determine if this was due to disorganization of the actin network, we measured Phalloidin fluorescence intensity as a function of distance along the Anterior-Posterior (AP) axis. We found that although there was an increase in the fluorescence intensity of the thin filaments, the periodicity was not affected by the expression of *Dystrophin* RNAi. (Fig. 6 A,B).

We then tested whether *Msp300* and *Dystrophin* had additive effects on microtubule organization. We first measured the propensity for colocalization of microtubules with F-actin and Dystrophin and found that in doubly heterozygous animals, similarly to controls, microtubules were more likely to colocalize with F-actin than Dystrophin (Fig. 6F). We next measured the intensity of tubulin immunofluorescence as a function of distance from the myonucleus in single and doubly heterozygote mutants. Both the *Msp300^compl^* heterozygotes and the *Dys^EP3397^/Msp300^compl^* double heterozygotes displayed less intense tubulin immunofluorescence peaks at the edges of the myonucleus indicating a reduced microtubule ring. Further, the decay in the tubulin immunofluorescence signal in both the *Msp300^compl^* heterozygotes and the *Dys^EP3397^/Msp300^compl^* double heterozygotes was reduced compared to controls (Fig. 6G). However, the profile of tubulin immunofluorescence intensity was similar in *Msp300^compl^* heterozygotes and the *Dys^EP3397^/Msp300^compl^* double heterozygotes which had distinct myonuclear spacing profiles. This suggested that the change in tubulin immunofluorescence intensity was not the underlying cause of disrupted myonuclear spacing. We then focused on the microtubules that were oriented on the dorsal-ventral axis of the myofibers, thus running orthogonally to the F-actin thin filaments, which would allow extended interactions between the microtubules and the sarcomeres. We found that the fraction of microtubules that colocalized with the actin thin filaments and extended on the dorsal-ventral axis was reduced in *Dys^EP3397^/Msp300^compl^* double heterozygotes compared to *Msp300^compl^* heterozygotes and *Dys^EP3397^* heterozygotes (Fig. 6H). In the *Msp300^compl^* heterozygotes and the *Dys^EP3397^* heterozygotes the proportion of dorsal-ventral microtubules colocalized with F-actin was reduced to 88% and 87% respectively. In the *Dys^EP3397^/Msp300^compl^* double heterozygotes only 79% of microtubule extending on the dorsal-ventral axis colocalized with F-actin.

Taken together, these data support a model whereby *Dystrophin* and *Msp300* regulate the maintenance of myonuclear spacing and microtubule organization in the fully differentiated larval myofiber. Furthermore, this work describes the first instance of a regulator of myonuclear spacing specific to the maintenance stage, with no role in the active movement of myonuclei in the developing myotube.

## Discussion

We have identified a new genetic interaction that maintains myonuclear spacing in contractile myofibers that is dependent on *Dystrophin* and *Msp300*. Although this is not the first work to implicate either of these factors in the spacing of nuclei in myofibers (Elhanany-Tamir et al., 2012; Wang et al., 2000), this is the first work to connect these two genes whose products show distinct localizations. Critically, we demonstrate that *Dystrophin* does not contribute to myonuclear spacing during embryonic development and contributes only to maintain myonuclear spacing in differentiated larval myofibers. These data are consistent with previous work that relied on genetic mutants to suggest that *Msp300* only contributed to myonuclear spacing during the later stages of development (Elhanany-Tamir et al., 2012; Wang et al., 2015). Additionally, Msp300, the *Drosophila* homolog of Nesprin-1G, and *Dystrophin* are linked to significantly different muscle diseases (Hoffman et al., 1987; Zhang et al., 2007). That they together regulate the spacing of myonuclei supports the growing evidence that nuclear position is a crucial feature of myofiber architecture and muscle function. Furthermore, the larval *Drosophila* myofibers are not regenerative. Thus, mispositioned nuclei are not an indication of ongoing repair. Therefore, maintaining the position of myonuclei is a core function of *Dystrophin* and *Msp300*.

We further show that the cell biological functions of *Dystrophin* and *Msp300* intersect in the regulation of microtubule organization. Although this is not the first work to implicate either *Msp300* or *Dystrophin* in the regulation of microtubule organization (Elhanany-Tamir et al., 2012; Khairallah et al., 2012), this work extends previous findings and specifically implicates their combined functions in the regulation of multiple features of microtubule organization. Specifically, we have identified the colocalization of microtubules with sarcomeres as an organizational feature that is lost in genotypes with the greatest disruption in myonuclear spacing. Microtubules that extend from the nucleus to the sarcomeres are ideally suited to maintain nuclear spacing during muscle contraction. Because the sarcomeres maintain a stable position during muscle contraction, they provide an anchoring point for the microtubules that extend from the nuclei and throughout the cytoplasm.

The mechanism by which *Dystrophin* regulates the colocalization of sarcomeres and microtubules requires further investigation but two hypotheses seem likely. The first posits that because *Dystrophin* and F-actin localization are primarily anti-correlated, *Dystrophin* excludes microtubules. The second posits that *Dystrophin* having both actin-binding domains and microtubule-binding domains crosslinks actin and microtubules. Previous biochemical analysis demonstrated that Dystrophin can directly alter microtubule dynamics by specifically increasing the time that microtubules spend in a paused state, neither growing or shrinking (Belanto et al., 2014). This finding, combined with our data indicating that the loss of *Dystrophin* correlated with decreased microtubule density supports the hypothesis that *Dystrophin* actively crosslinks actin and microtubules.

The mechanism by which *Dystrophin* and *Msp300* cooperate to regulate microtubule organization is likely indirect. The primary localizations of Dystrophin and Msp300 are at the sarcolemma and the outer nuclear membrane respectively. However, Msp300 has been shown to have cytoplasmic localizations (Elhanany-Tamir et al., 2012), which could enable biochemical interactions with Dystrophin. Yet, the data presented here does not necessitate a biochemical interaction and is instead consistent with an indirect interaction via the microtubule network. One possible mechanism for the functional interaction between *Msp300* and *Dystrophin* is separate effects on microtubules that are each required to maintain nuclear spacing. Specifically, *Msp300* may be necessary for the proper attachment of microtubules to myonuclei. This is supported by data in this work and by data in previous work (Elhanany-Tamir et al., 2012; Wang et al., 2015). Alternatively, *Dystrophin* is necessary for colocalization of microtubules and sarcomeres. A simultaneous reduction in the number of microtubules attached to myonuclei and a reduction in the number of microtubules that colocalize with sarcomeres may together dramatically reduce the number of microtubules that extend from the myonucleus to the sarcomeres to stabilize myonuclear spacing.

In total, the data presented here demonstrate a functional link between two genes, *Msp300* and *Dystrophin*, both of which are mutated in patients with disparate muscle diseases. The functional interaction is manifest in two different cellular phenotypes, disruptions in microtubule organization and disruptions in myonuclear spacing. Crucially, these two cellular disruptions are common phenotypes associated with disparate muscle disorders supporting the hypothesis that each are critical contributors to muscle health.

## Materials and Methods

### Drosophila genetics

All stocks were grown under standard condition at 25°C. Stocks used were *apRed* which expresses *DsRed* fused to a nuclear localization signal downstream of the *apterous* mesodermal enhancer allowing for visualization of nuclei within the LT muscles during embryonic stages (Richardson et al., 2007), *bocks^DP01391^* (21846; Bloomington *Drosophila* Stock Center), *klar^1^* (3256; Bloomington *Drosophila* Stock Center), *Msp300^compl^* (Courtesy of T. Volk: Elhanany-Tamir et al., 2012) *koi^EY03560^* (20000; Bloomington *Drosophila* Stock Center), *ote^DB^* (5092; Bloomington *Drosophila* Stock Center), *ote^B279^* (16189, Bloomington *Drosophila* Stock Center), *Dys^EP3397^* (17121; Bloomington *Drosophila* Stock Center), *Dys^E17^* (63047; Bloomington *Drosophila* Stock Center), *Khc^8^* (1607; Bloomington *Drosophila* Stock Center), *Dhc64C^4-19^* (5274; Bloomington *Drosophila* Stock Center), *ens^swo^* (Metzger et al., 2012), UAS-mCherry RNAi (35785; Bloomington *Drosophila* Stock Center), UAS-Dys RNAi (55641; Bloomington *Drosophila* Stock Center), UAS-Shot RNAi, UAS-Dhc RNAi, UAS-Scgα RNAi (34027; Bloomington *Drosophila* Stock Center), UAS-Dyb RNAi (36101; Bloomington *Drosophila* Stock Center), UAS-Dg RNAi(34895; Bloomington *Drosophila* Stock Center), UAS-Syn1 RNAi (27504; Bloomington *Drosophila* Stock Center) and UAS-Klar.3’ΔS (25103; Bloomington *Drosophila* Stock Center). Mutants were balanced and identified using *IF*/*CyODGY* for the second chromosome and *Ly, hs-hid*/*TM6DGY* for the third chromosome. The exception to this is for Msp300^compl^ which is balanced over *CyOGFP*. All UAS expression was driven specifically in larval muscle using *DMef2-GAL4*, *apRed* or *MHC-GAL4*. Controls were *twist-GAL4*, *apRed, MHC-GAL4,* and *Dmef2-GAL4*, *apRed Drosophila* lines as indicated in figure legends. Third chromosome mutant alleles had *twist-GAL4*, *apRed* on the second chromosome and second chromosome mutant alleles had *Dmef2-GAL4*, *apRed* on the third chromosome. The *twist-GAL4*, *apRed* and *Dmef2-GAL4*, *apRed Drosophila* lines were made by recombining the *apRed* transgene and the specific GAL4 driver.

### Sample Preparation & Immunohistochemistry

Embryos were collected at 25°C and then dechorionated by submersion in 50% bleach for 5 min. Embryos were then washed with water and then fixed in 10% formalin (HT501128; Sigma-Aldrich) diluted 1:1 with heptane and placed on an orbital shaker that rotated at a rate of 300 rpm for 20 min. Embryos were devitellinized by vortexing in a 1:1 methanol:heptane solution. Embryos were stored in methanol at −20°C until immunostaining. Larvae were dissected as previously described (Mandigo et al., 2019)

Antibodies for embryo staining were used at the following final dilutions: rabbit anti-dsRed, 1:400 (632496; Clontech); rat anti-tropomyosin, 1:200 (ab50567; Abcam). Antibodies for larval staining were used at the following final dilutions: Rabbit anti-Dystrophin, 1:150 (ab15277; Abcam), Rabbit anti-αTubulin, 1:200 (T6199, Sigma-Aldrich), and mouse anti-αTubulin, 1:200 (T6199, Sigma-Aldrich). Conjugated fluorescent secondary antibodies used for embryo and larval staining were Alexa Fluor 488 donkey anti-mouse-IgG (1:200), Alexa Fluor 647 donkey anti-mouse-IgG (1:200), Alexa Fluor 647 donkey anti-rabbit-IgG (1:200), and Alexa Fluor 647 donkey anti-mouse-IgG (1:200; all Life Technologies). Acti-stain 555-conjugated phalloidin (1:400; PHDH1-A; Cytoskeleton) and Hoechst 33342 (1 μg/ml; H3570; Life Technologies) were used for larval staining only. Embryos and larvae were mounted in ProLong Gold (P36930; Life Technologies) and imaged with an Apochromat 40×/1.3 numerical aperture (NA) objective with a 1.0× optical zoom for all embryo and larval images on a Zeiss 880 LSM. Additionally, larval images were acquired with bidirectional tile scan with an online stitching threshold of 0.70.

### Analysis of nuclear position in embryos

The position of nuclei was measured in stage 16 embryos, which is the latest stage before cuticle development blocks the ability to perform immunofluorescence microscopy. Embryos were staged primarily by gut morphology as previously described (Folker et al., 2012). Images were processed as maximum intensity projections of confocal *z*-stacks using FIJI. The positioning of the nuclei was measured using the segmented line function in FIJI to determine the distance between either the dorsal end of the muscle and the nearest nucleus or the ventral end of the muscle and the nearest nucleus. All four LT muscles were measured in four hemisegments from each embryo. At least 10 embryos from each of two independent experiments were measured for each genotype. Statistical analysis was performed with Prism 6.0. Student’s *t*-test was used to assess the statistical significance of differences in measurements between experimental genotypes and controls.

The nuclear distribution ratio was calculated by dividing the area occupied by the dorsal nuclear cluster by the area occupied by ventral nuclear cluster. Statistical analysis was performed with Prism 6.0. Student’s *t*-test was used to assess the statistical significance of differences in measurements between experimental genotypes and controls.

### Analysis of myonuclear position in larvae

We measured myonuclear position in larvae with as previously described (Auld et al., 2018; Collins & Mandigo et al., 2017). First, the area and length of the muscle were measured. Next, the position and number of nuclei were calculated using the multipoint tool in FIJI to place a point in the center of each nucleus. The position of each nucleus was used to calculate the actual internuclear distance. The maximal internuclear distance was then determined by taking the square root of the muscle area divided by the nuclear number. This value represented the distance between nuclei if their internuclear distance was fully maximized. The ratio between the actual internuclear distance and the maximal internuclear distance was then used to determine how evenly nuclei were distributed. This method normalized the internuclear distance to both the nuclear count and the muscle area, which resulted in a more representative means of comparison between muscles, larvae, and genotypes. A total of 24 ventral longitudinal (VL3) muscles were measured from at least three larvae with at least three VL3 muscles measured from each larva from at least two biological replicates. Statistical analysis was performed with Prism 6.0. Student’s *t*-test or ANOVA with multiple comparisons depending on number of genotypes used.

### Airyscan Acquisition & Analysis of Microtubule Organization in Larvae

These two analyses utilized high-resolution images taken using Elyra Airyscan processing. Larval dissections were prepared as previously described and imaged on the same microscope and objective as described above with a 3.0× optical zoom. All step sizes and speed settings were set to ‘optimal’ within the Zen Black acquisition software for every image. Elyra Airyscan post processing was done using Zen Blue.

The first method used a code which automatically detected nuclei within a myofiber and drew concentric circles of increasing radii from the center of the nucleus. The radius increased by 2.5μm per ring. The intensity of signal within each ring was measured and plotted as a function of circle radius. Concentric circle analysis was run on five non-clustered nuclei from three different myofibers across two animals. Statistical analysis was performed with Prism 6.0. Student’s *t*-test was used to assess the statistical significance of differences in measurements between experimental genotypes and controls.

The second method used to measure microtubule organization was co-localization of microtubules with actin or dystrophin bands. Regions of interest were defined in FIJI based on areas of strong microtubule banding that were not directly associated with the microtubule ring around nuclei. At least 2 regions of interest were selected from different myofibers. All myofibers came from different animals. Pearson’s correlation coefficients for MTs with actin and MTs with dystrophin were determined separately using the CoLocalization plugin for FIJI. Statistical analysis was performed with Prism 6.0. Student’s *t*-test was used to assess the statistical significance of differences in measurements between experimental genotypes and controls.

Alignment of microtubules was determined by tracing ROIs along all sections of microtubules that were oriented in the dorsal-ventral direction Using FIJI. Then, the traced ROIs were placed over the actin channel of the same image and scored as either overlapping and tracing, or no-overlap/overlap but not aligned with the actin-based thin filaments.

Myonuclei without rings were measured by subtracting the number of rings in the microtubule channel of a myofiber from the total number of myonuclei seen in that same myofiber via Hoescht staining.

### Analysis of actin and dystrophin localization

Images of whole myofibers were split into individual channels and a line was drawn perpendicular to the dorsal:ventral axis and parallel to the anterior:posterior axis of the myofiber in both the actin and dystrophin channels. The size and position of the line was maintained in both images via the ROI manager in FIJI. Then, intensity signals based on position were measured and subsequently plotted in excel to show peak alignment/misalignment of both channels.

## Acknowledgements

We thank Bret Judson and the Boston College imaging core for infrastructure and support with microscopy. We thank the Bloomington *Drosophila* Stock Center for stocks that were used in this study (NIH P400D018537). We thank Talila Volk’s group at the Weissman Institute for providing *Msp300* null allele stocks.

---

**Supplementary Figure 1:**
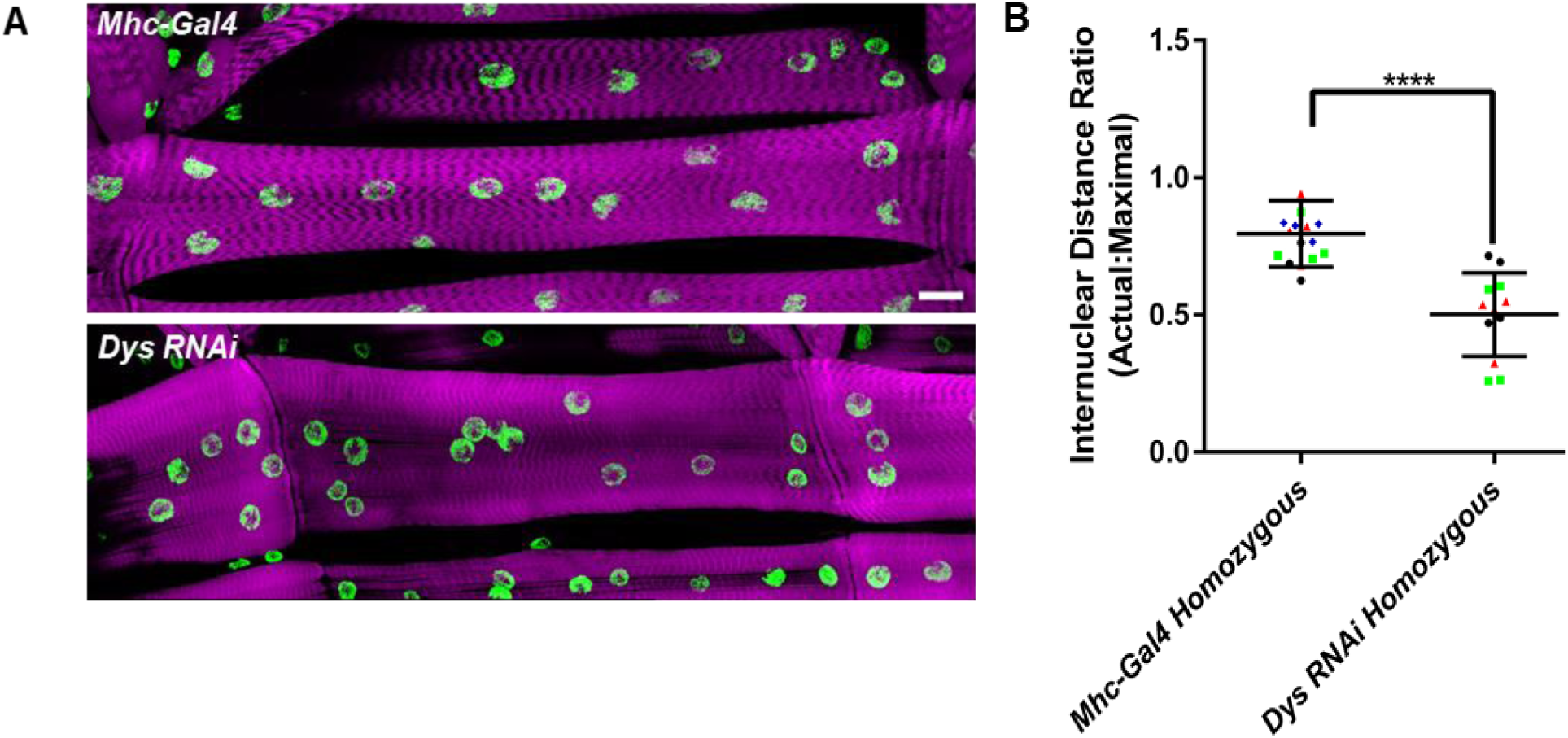
Mhc-Gal4 homozygotes do not have disrupted spacing when compared to animals homozygous for RNAi insertions at *attp40*. **(A)** Fluorescence images of VL3 muscles with indicated genotypes. Magenta, phalloidin/muscles; green, Hoechst/nuclei. Scale bar = 25μm. **(B)** Internuclear distance ratio in larval muscles from animals with the indicated genotype. Each point is one myofiber. Myofibers from the same animal are the same shape and color. Data compared by t-test. Four myofibers were analyzed from four animals. P < 0.0001

